# A *Drosophila* larvae-inspired vacuum-actuated soft robot

**DOI:** 10.1101/2022.05.08.491074

**Authors:** Xiyang Sun, Akinao Nose, Hiroshi Kohsaka

## Abstract

Peristalsis is one of the most common locomotion patterns in limbless animals. This motion is generated by propagating muscular contraction and relaxation along the body axis. While the kinematics of peristalsis has been examined intensively, the kinetics and mechanical control of peristalsis remain unclear, partially due to the lack of suitable physical models to analyse the force and temporal control in soft-bodied animals’ locomotion. Here, based on a soft-bodied animal, *Drosophila* larvae, we proposed a vacuum-actuated soft robot replicating their crawling behaviour. The soft structure, made with hyperelastic silicon rubber, was designed to mimic the larval hydrostatic structure. To estimate the adequate range of pressures and time scales for control of the soft robots, a numerical simulation by the finite element method was conducted. Pulse-Width-Modulation (PWM) was used to generate time-series signals to control the vacuum pressure in each segment. Based on this control system, the soft robots could exhibit the peristaltic pattern resembling fly larval crawling. The soft robots reproduced two previous experimental results on fly larvae: slower crawling speed in backward crawling than in forward crawling, and the involvement of segmental contraction duration and intersegmental delay in crawling speed. Furthermore, the soft robot provided a novel prediction that the larger the contraction force, the faster the crawling speed. These observations indicate that the use of soft robots could serve to examine the kinetics and mechanical regulation of crawling behaviour in soft-bodied animals.

## 1. Introduction

Over the past few decades, robotics researchers have been drawing inspiration from diverse species of animals to design robots [1, 2]. A recent approach to building animal-inspired robots is utilizing soft materials to construct flexible structures, mainly because the flexibility enables adaptive motions [2, 3]. Furthermore, the development of soft robots has provided insights into the biological mechanisms of animal motion [4, 5]. In particular, soft robots are useful for understanding the kinematics of soft-bodied animals’ behaviours since their locomotion has a high degree of freedom. The use of physical simulation is indispensable in examining the complicated dynamics [6]. The development of biomimetic soft robots has provided valuable platforms for both robotics and neuroscience research fields by referring to animals [7], including caterpillars [1], earthworms [8], and octopi [9].

Crawling behaviour is generated by the propagation of segmental contraction along the body axis to move the body forward [10]. Crawling is one of the basic animal motions used to move the body in one direction and has been mimicked by soft robots, including worm-like robots [8], hornworm-like robots [1], snake-like robots [11], and multigait soft robots [12]. Previously, flexible braided mesh-tube structures, Meshworm and FabricWorm, were designed based on the antagonistic muscular arrangement of earthworms [8, 13] and were capable of exhibiting crawling. They used shape-memory alloys and linear springs as actuators, respectively. Despite these recent advances in soft robots for studying crawling behaviour, how crawling properties, including speed, are realized and regulated remains unclear [14, 15].

Larvae of fruit flies, *Drosophila melanogaster*, have provided an excellent model of a soft-bodied organism to investigate peristaltic mechanisms due to their relatively simple structure, stereotyped behaviours, and accumulated knowledge of their neural circuits [16, 17]. The third instar fly larva is about 4 mm long and has a segmented body. The dominant larval behaviour is forward crawling, which involves the sequential translation of body segments via muscle contraction along the body axis. However, the larvae also exhibit backward behaviour, with the same muscles activated in the opposite sequence [18]. One soft maggot robot was previously designed to mimic larval muscular organization and replicate larval crawling [19]. It consisted of a series of pneumatic chambers that enabled body deformation by expansion. Although coordinated motions were generated, this previous maggot robot did not realize crawling behaviour.

Pneumatic circuits have been used as an actuator in soft robots. By increasing and decreasing internal pressure in chambers of soft material, adaptive and versatile motion can be realized [14]. There have been various applications of pneumatic circuits to soft robots to perform multiple gaits [14], a hybrid of hard and soft robots [20], and gloves for hand rehabilitation [20].

In this work, we propose a new soft robot that can mimic larval crawling through reference to the properties of fly larvae. To mimic the contraction of body segments, a vacuum source and solenoid valves were used for actuation control. We implemented an asymmetric interface between the robot and a ground substrate to enhance the forward crawling rather than backward crawling. This larval robot successfully exhibited a crawling motion. Perturbation experiments suggest that the contraction force and segmental phase delay are critical for the crawling speed. This study indicates that our vacuum-based soft robots could contribute to a better understanding of the mechanisms of crawling behaviour and the development strategy of soft robots with faster crawling speeds.

## 2. Methods

By mimicking the biological properties of fly larvae, we proposed a soft robot consisting of the following three components: A) a body structure with a chain of elliptical cylinder-like segments, B) a vacuum-actuated control system, and C) software for controlling and monitoring the motion of the maggot-like soft robot.

### 2.1 Body structure

When designing our soft larval robot, three structural properties in fly larvae were taken into account: 1) hydrostatic structure, 2) repetitive muscular patterning and 3) asymmetric substrate interaction via denticle bands.

First, the fly larval body is filled with body fluid, and the internal pressure and the tension of the body wall play a role in supporting their body shape. To replicate the larval hydrostatic properties, we adopted a pneumatic structure with silicone rubber among the current soft actuator candidates [2, 20]. Second, the configuration of muscles in the body wall is segmentally repeated in fly larvae. The larval body consists of 11 segments: three thoracic segments and eight abdominal segments. Although the terminal segments (the first thoracic (T1) and the last abdominal (A8) segments) have specialized structures, the other nine segments have an almost consistent structure. The larval length and width were examined as 3.69±0.56mm and 0.66±0.09mm [21]. The length-to-width ratio of a single segment in the major middle segments is about 0.5 (Supplementary Figure 1). To mimic the larval shape, the individual segment was designed as an elliptical cylinder chamber with a flat plane at the bottom. We constructed two different soft robots with different segmental length-width ratios: The width of all the segments was 30 mm, whereas the axial length of the segments of the three robots was 20 mm and 30 mm, respectively. Accordingly, their length-to-width ratios of them were 2/3 and 1, respectively. To simplify the robot while allowing us to analyze the propagation of segmental contraction, we set the number of the segments in one robot as five (Figure 1). Third, spike-like structures align segmentally at the bottom of the fly larvae, named denticle bands. Denticle bands act as anchorage points to the ground during the propagation of segmental deformation [22]. Each denticle band normally consists of six rows of denticles in a larva. The hooked tips of four out of the six rows point posteriorly while the remaining two rows point anteriorly [22], suggesting that the friction between the ventral body surface and the ground substrate should be different between forward and backward motions. To mimic this asymmetric denticle structure between anterior and posterior directions, we implemented an asymmetric friction structure by glueing a piece of paper (Whatman paper 1001-917) at the anterior side of the segment boundary (shown as the thick lines in Supplementary Figure 1). Silicone rubber is stickier than paper; hence the physical contact between the silicone rubber and a ground substrate would generate larger friction than the one between paper and the ground substrate. By virtue of this property and the geometric fact that the angle of the segment boundary depends on the direction of the propagation of segment contraction, asymmetric friction during locomotion in different directions could be realized.

**Figure 1.**
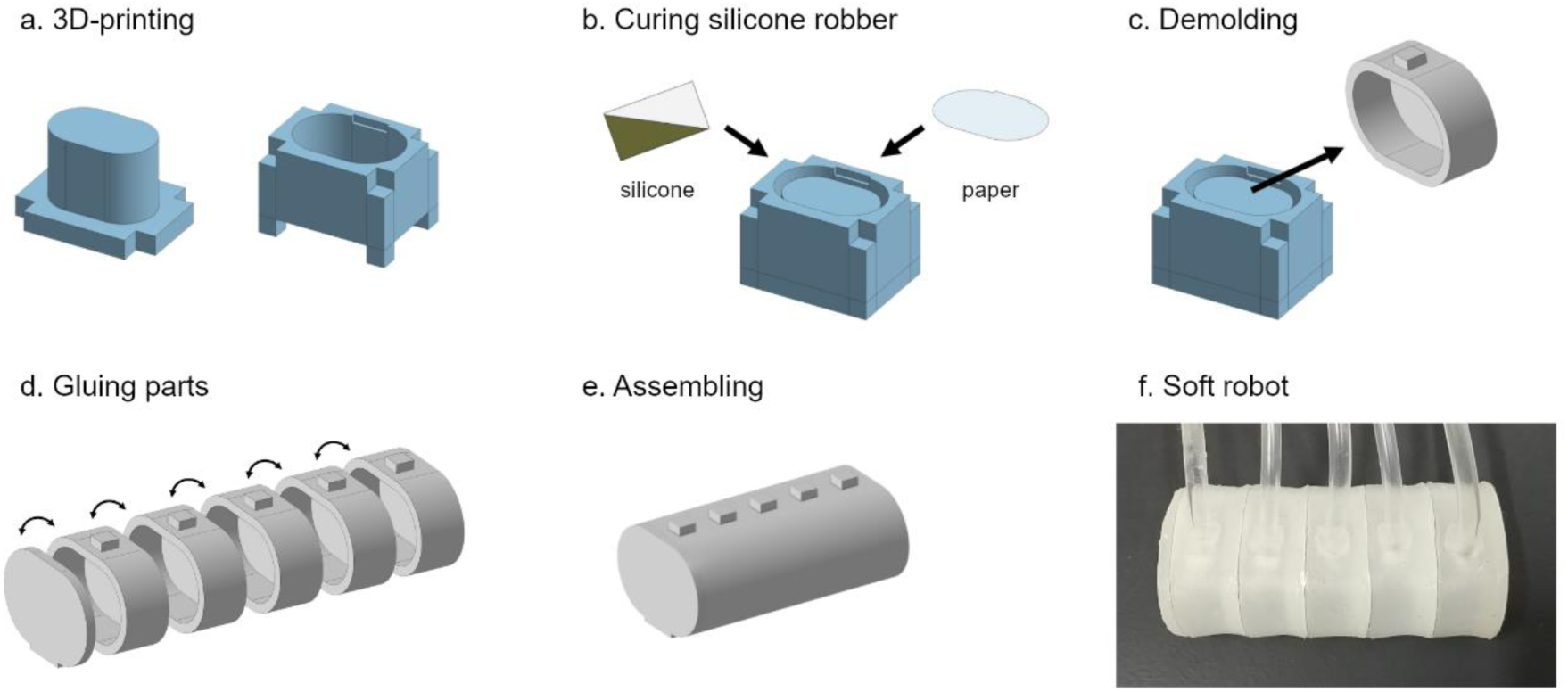
The fabrication process for the soft robot.

We fabricated soft robots based on the design shown above. First, the moulds for the soft robots were printed with Acrylonitrile Butadiene Styrene (ABS) material with a 3D printer (WANHAO Duplicator 4S). Then, the moulds were integrated (Figure 1A), and silicone rubber (Ecoflex 00-30) was poured into the space between the inner and the outer moulds. After the gap was filled up with silicone rubber, a piece of paper (Whatman paper 1001-917) was inserted between the anterior surface of the liquid rubber and the mould (Figure 1B). The rubber was cured in a drying oven (at 80°C for about 20 minutes) and cooled down at room temperature. Then, the single segment structure was demolded, as shown in Figure 1C. Finally, five segmental structures were glued together using the Ecoflex mixture to build the final structure (Figures 1D and 1E). An example is shown in Figure 1F.

### 2.2 Vacuum control system

In order to enable the segmental chambers to contract like body segments in fly larvae, we took advantage of a vacuum-based actuator. We built a system consisting of pneumatic circuits, electrical circuits (dashed and solid lines respectively in Figure 2), and a graphical user interface software (implemented on “Computer” in Figure 2).

**Figure 2.**
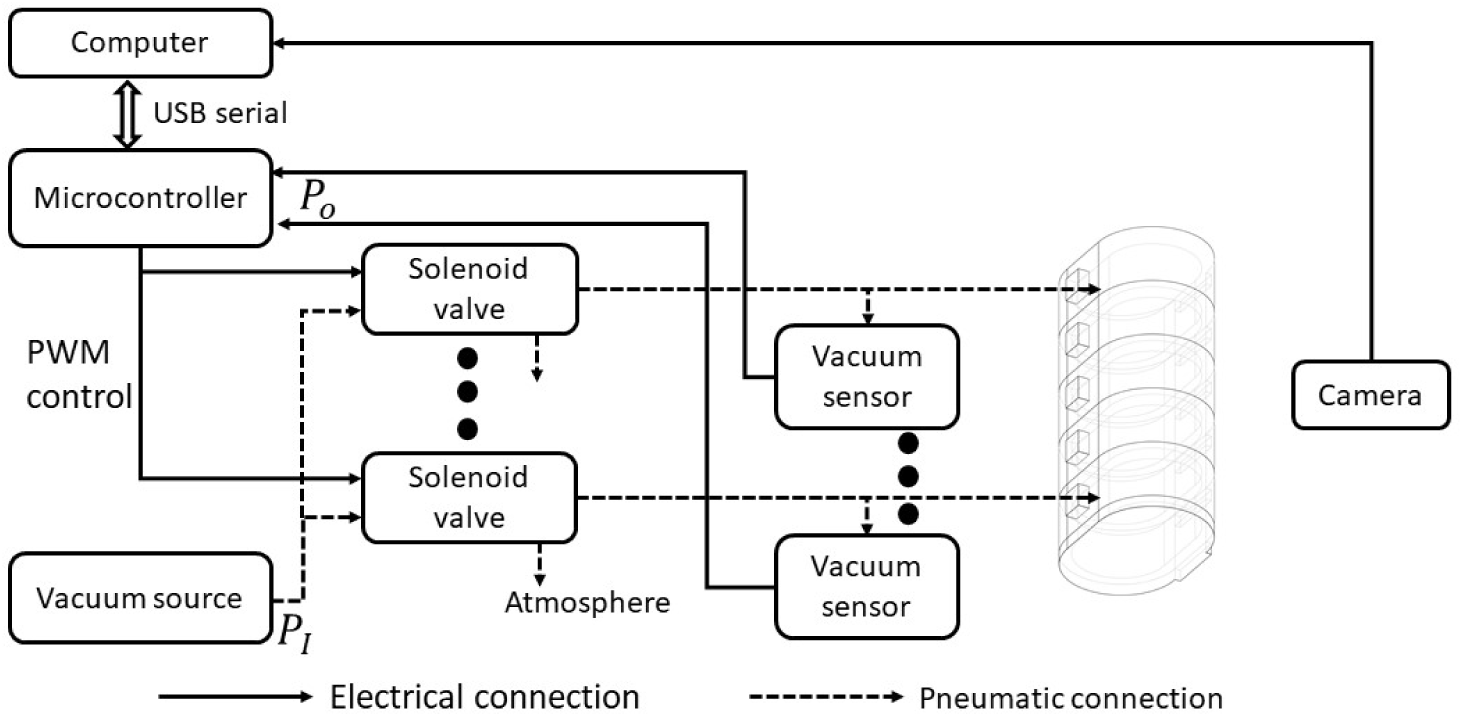
The framework of the whole system.

The pneumatic circuits were established to control the pressure in each segment chamber. Different from previous larvae-like robots based on segmental expansion [19], in this study, we adopted contraction as a driving force for larval locomotion to mimic muscular contraction in fly larvae. Pneumatic pathways linked a vacuum source (TAITEC VC-15s, with a pressure range from -110kPa to 0kPa), three-way solenoid valves (ZHV 0519), vacuum pressure sensors (MPXV6115V), and robotic chambers (see section 2.1). Each chamber and the vacuum source were connected through a vinyl tube and a solenoid valve. The tube was flexible and lightweight (1.4 g) compared with the soft robots (16.9 – 29.9 g). Even when we held the tubes, the motion of the soft robots was not disturbed, showing that the tube has little effect on the kinematics of the soft robot. The pressure within each chamber was regulated by gating the solenoid valves.

Electrical circuits were built to control the pressure valves for individual chambers. A microcontroller (Arduino Mega) was selected considering the number of PWM pins and the output voltage. The microcontroller was connected to solenoid valves and vacuum pressure sensors to regulate and monitor robot deformation. The microcontroller’s detailed electrical circuits are presented in Supplementary Figure 2, including modules for the solenoid valves and vacuum sensors. Here, an NPN transistor (TIP120) modulates an external high-power source to drive the solenoid valves based on the PWM signal from the microcontroller. The 6V external power, provided via a DC-DC power supply regulator (ARD-PWR), was the maximal operational voltage for the solenoid valves. Since the valves possessed non-negligible inductance, diodes (1N4007) are used in a flyback configuration to prevent a large back-electromotive force (back emf), which could damage these valves. These diodes could dissipate the remaining energy. The vacuum pressure, in the range of –115 to 0 kPa relative to the standard atmosphere, was measured by the output of the voltage-based vacuum sensor based on the following equation:

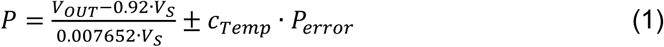

where *P* is the vacuum pressure, and *V*_*OUT*_ and *V*_*S*_ represent the output voltage and voltage supply, respectively, in the pressure sensor. *P*_*error*_ is the pressure error, which indicates the measurement error in the pressure at a standard temperature (the dimension of *P*_*error*_ is pressure). *c*_*Temp*_ is a temperature factor for the pressure error, which is 1 within the temperature ranges from 0 to 85°C (*c*_*Temp*_ is dimensionless). According to the sensor datasheet, the maximum value of *P*_*error*_ is 1.725 kPa within the range of our experiments. Low-pass filters were applied to pressure signals from vacuum sensors (Supplementary Figure 2).

### 2.3 Software for controlling and monitoring soft robot locomotion

To monitor the position of the segment boundaries and their segmental pressures in operation in real-time, we designed a graphical user interface (GUI) (Supplementary Figure 3) based on Python libraries (Tkinter and pySerial). The USB camera (HOZAN camera with 12mm lens) is attached to a bracket right above the soft robot. Its frame rate is up to 40 frames per second, used in the following experiments. The images were sent to the computer via a USB connector, and the robotic motion could be observed on the GUI (Figure 3 right, Supplementary Figure 3). The position of the front end (“head”) and rear end (“tail”) was measured by a ruler shown in Figure 3. Meanwhile, pressure signals of robot segments were delivered from the Arduino board to the computer. Example locomotion and pressure signals are shown in Supplementary Figure 3.

**Figure 3.**
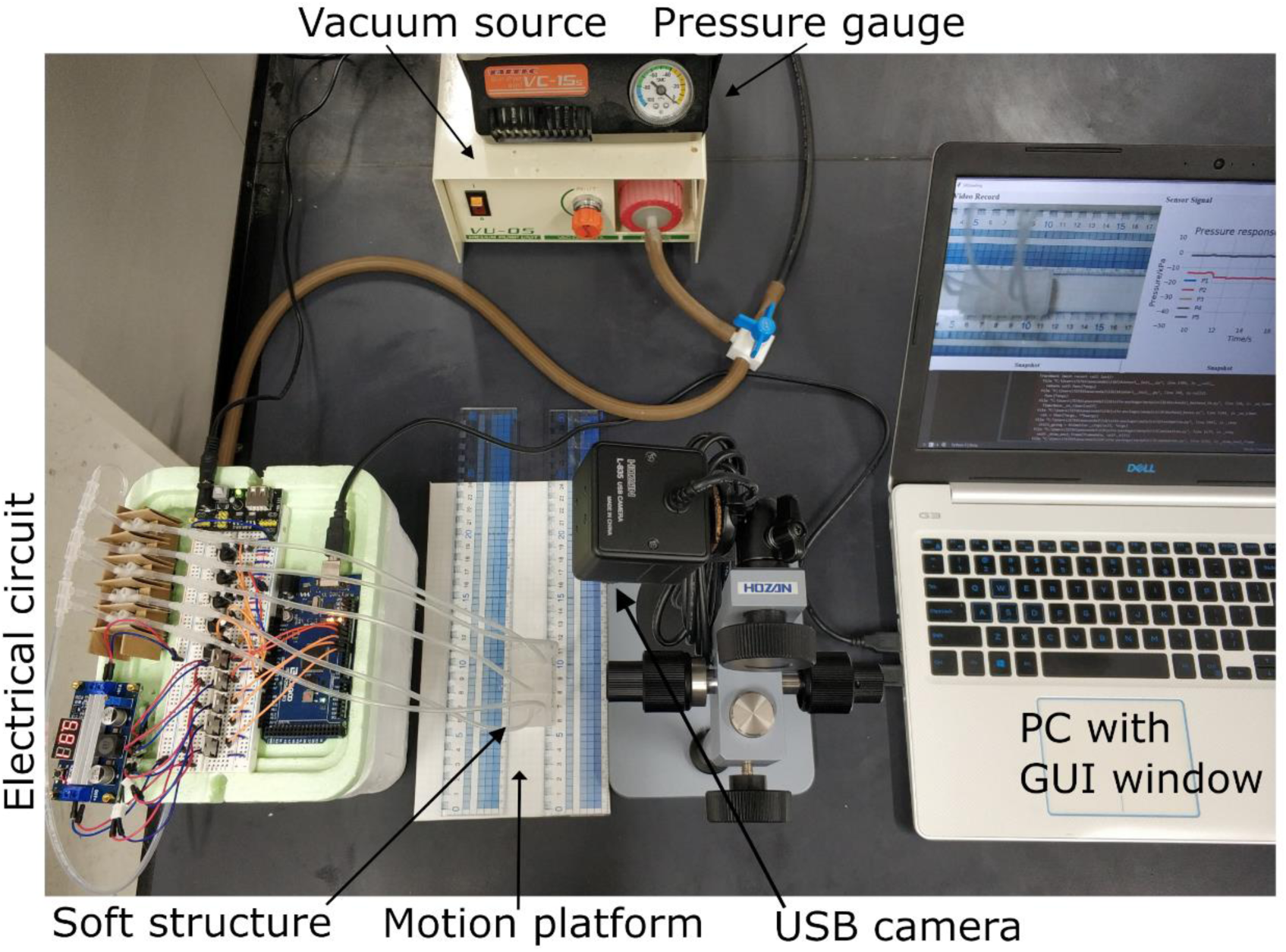
System overview.

## 3. Simulation

To determine the suitable ranges of the pressure and time scale for robot deformation, we applied the finite element method (FEM) to simulate our soft robot via the commercial FEM software (Abaqus). To achieve a reliable simulation, suitable viscoelastic physical models were required. In our project, the main soft body was constructed using hyperelastic material (Ecoflex 00-30). In line with previous modelling and validation work [6], the Ogden model was adopted to model this material. The strain energy potential function *U*, which describes the elastic properties of the material, is defined as:

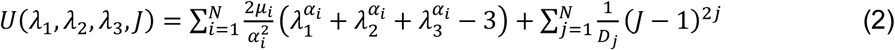

where *μ*_*i*_ and *α*_*i*_ (*i* = 1, 2, 3) are the primary fitting parameters, and *λ*_*j*_ (*j* = 1, 2, 3) are the stretches along the x, y, and z axes. The material constant *N* was set as three. The second summation term contains fitting parameters *D*_*j*_ (*j* = 1, 2, 3) to the volumetric deformation and the material Jacobian matrix *J*. We referred to the previous parameters for this hyperelastic material [6]. Meanwhile, regarding the paper (Whatman paper) on the intersection of soft robots, the material density is 4.83e-10*t*/*mm*^3^, and its young’s module is 1.71GPa and poisson ratio is -0.3 [23]. Considering the segmental deformation under vacuum pressure, we also configured the self-contact condition for the interaction step.

We calculated the responses of the soft robot under distinct pressures using the FEM software Abaqus (Figure 4). With respect to the nonlinear dynamics within our model, we configured parameters for incrementation to ensure a successful simulation: the maximum number of increments as 100 and the minimum increment size as 1e-15. Here, the example soft robot with five 20 mm segments was indicated as 20mmx5-SR (20 mm, five segments, and Soft Robot), and a similar shorthand notation was used for 30 mm segments 30mmx5-SR. Two points at the bottom of the terminal segment (marked by red points in Figure 4A) were used to monitor the head and tail segmental position in the longitudinal axis to record their asymmetric deformation. When the negative pressure of -10kPa was applied to the most anterior chamber (head chamber) of 20mmx5-SR, the head marker exhibited a negative displacement (Figure 4C), which indicated that the head segment was contracted and the end of the head moved backwards. The head marker moved faster when a larger absolute value of the negative pressure (from -20kPa to -70kPa) was applied to the head chamber. This observation indicated that the kinematics of the segment dynamics could be regulated by the pressure within the soft robot chambers. Since there was no asymmetricity along the body axis in the FEM simulation, when negative pressures were applied at the most posterior chamber (tail chamber), the tail marker showed similar displacements but in the opposite direction (Figure 4F). The larger soft robot (30mmx5-SR) exhibited faster segment contraction (Figure 4D and 4G). In either case, the movement was carried out in less than 1 second, comparable to the stride duration in fly larval locomotion.

**Figure 4.**
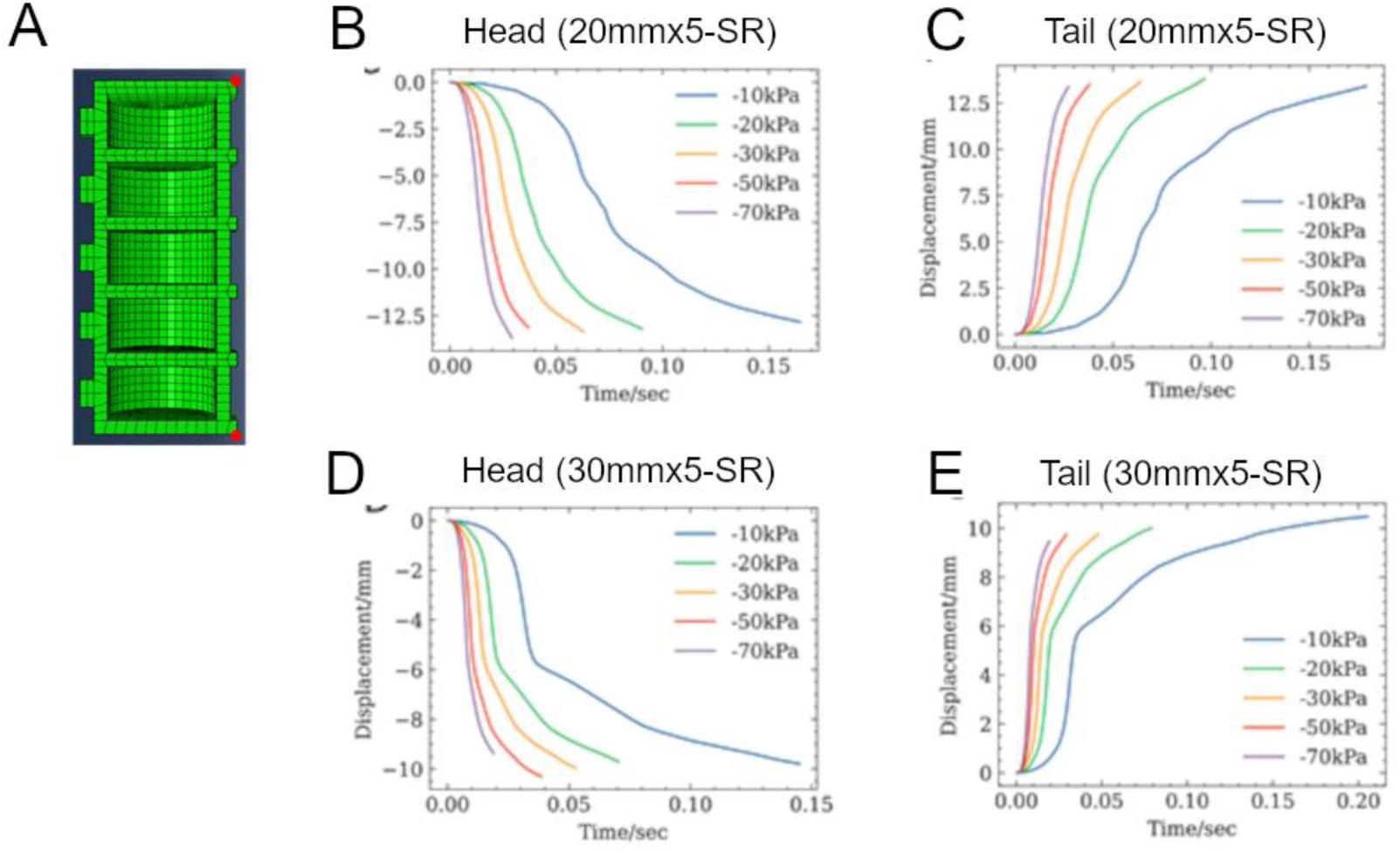
Simulation of the soft robots by FEM.

## 4. Results

### 4.1 Test for control signal and Comparison with simulation responses

In this study, we analyzed the locomotion of the soft robots under conditions with varied contraction forces and distinct intersegmental phases. Since the pressure of the vacuum source was constant, we tried to realise varied pressures by controlling the duty cycle of the gating of solenoid valves. To this aim, we adopted the Pulse-Width-Modulation (PWM) method [11] to control the temporal patterns of the chamber pressures. In this method, a series of pulses are generated, and the frequency and the duty cycle of the pulses can be tuned. We set the frequency as 1000 Hz because it was fast enough to reproduce fly larval crawling that occurred on the scale of 100 ms. The frequency of 1000 Hz was realized by setting a parameter OCRnA for the Arduino Mega board as 249 (= 16 MHz / 1000 Hz / 64 – 1). On the other hand, the duty cycle could be adjusted by OCRnB/OCRnC, another parameter for the board. Both OCRnA and OCRnB/OCRnC are in the range from 0 to 255 (Figure 5A). OCRnB/OCRnC set the value thresholding of a sawtooth timer signal to produce PWM signals with varied duty cycles (Figure 5A). We then used the resulting PWM signal to control segmental deformation via vacuum pressure. The deformation of the head segment was monitored under different duty cycles (Figure 5B). The results showed that the segmental deformation couldn’t be observed when the OCRnB is smaller than 190, which means that the solenoid valve doesn’t work in this case. On the other hand, when OCRnB was 195 or more, the contraction of the head segment was observed. In particular, as OCRnB increased, the speed of the deformation increased, which suggested that the contraction force within the head segment chamber was higher when the duty cycle was larger. Accordingly, we succeeded in temporally controlling the pressures within the segments of the soft robot. Even when we changed the waveform from the square to others, such as sinusoidal and saw-like waveforms, the temporal profiles of chamber pressure were similar to that with a square waveform input (Supplementary Figure 4). Then we decided to use the square waveform to control the chamber pressure in the following analyses.

**Figure 5.**
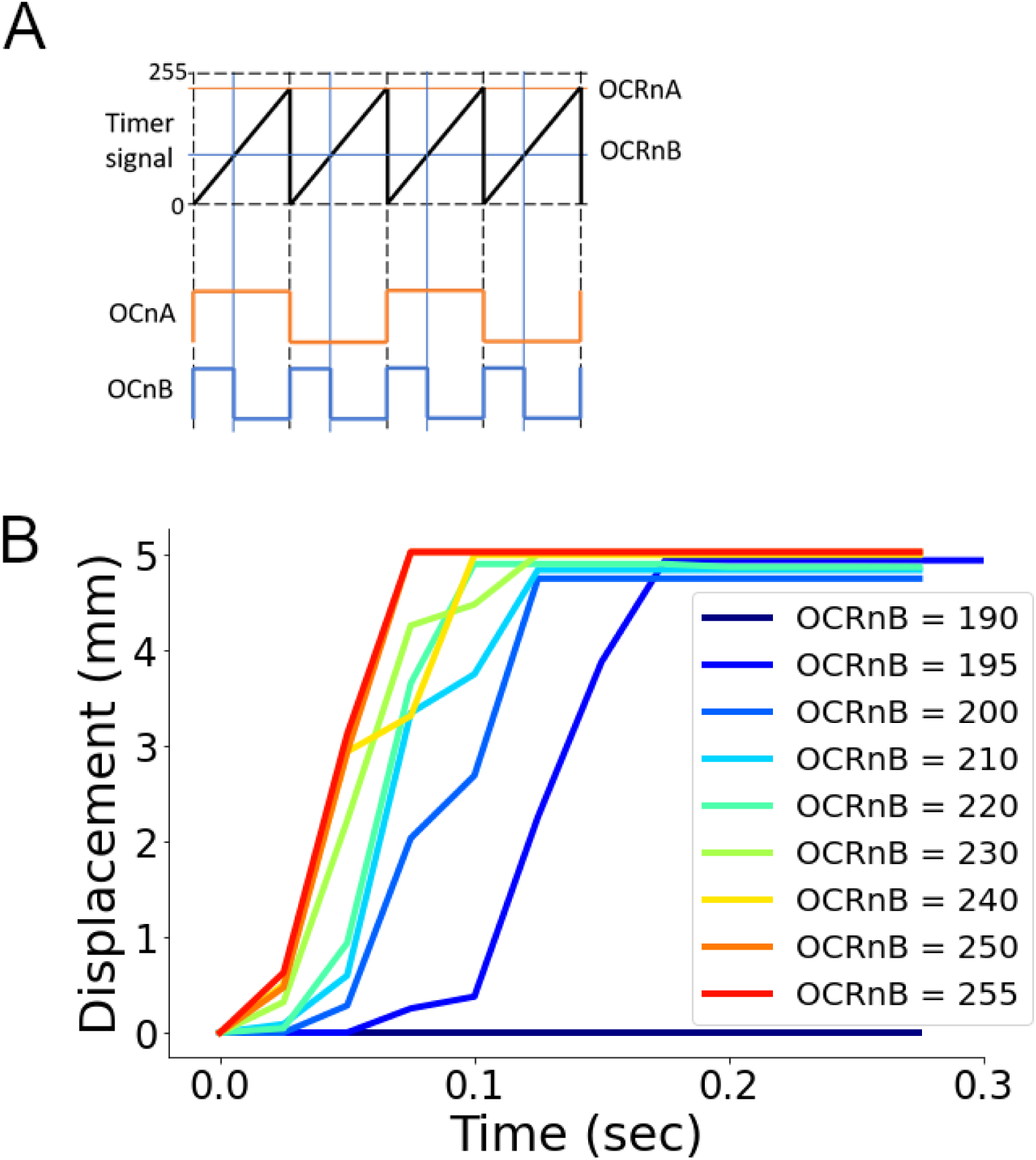
Test for a suitable range in the PWM control.

Referring to the results from the FEM simulation (see section 3), we analyzed the response of the soft robots to the pneumatic operation (Figure 6). When negative pressure was applied to the head chamber, the head marker on the 20mmx5-SR exhibited negative displacements, which was consistent with the FEM simulation results (Figure 6A). The displacement gradually reached a stable value in less than 1 second. A similar tendency was observed when negative pressures were applied to the tail chamber (Figure 6B). The larger soft robot (30mmx5-SR) showed weaker displacement than 20mmx5-SR (Figures 6C and 6D). All of these observations in the measurement in the soft robots were consistent with the FEM results. These analyses indicated the larger mass of soft robots, such as 30mmx5-SR, would make the robot harder to deform and move. Hence, we described experimental results for the 20mmx5-SR soft robot in the following sections.

**Figure 6.**
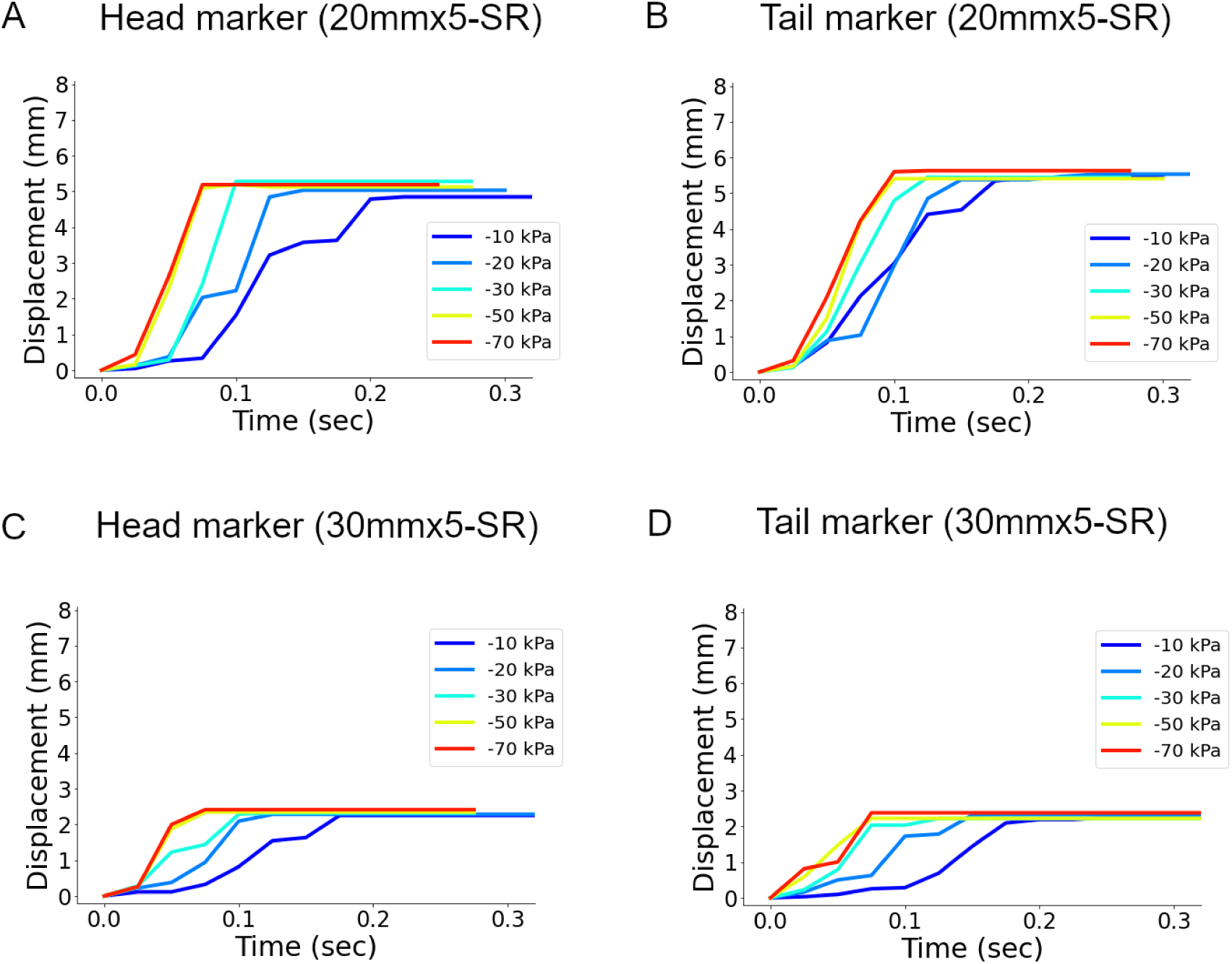
Segmental deformation in the soft robots.

One striking difference between the simulation and the soft robot experiment was the existence of asymmetric friction. As described in Supplementary Figure 1C, the friction at the interface between the soft robot and the ground substrate was asymmetric. Due to this effect, the tail marker showed a larger displacement upon applying negative pressure to the tail chamber than the head marker did upon the negative pressure applied to the head chamber. (Figure 6).

In summary, we implemented and optimized three properties in the soft robot to mimic fly larvae: The contraction of chambers instead of the expansion, which was used previously [19], the asymmetric feature of the interface between the soft robot and the ground substrate, and optimal ranges in pressure, time scale, and the size of the chambers.

### 4.2 Robotic locomotion and its quantification

We tested the crawling ability of the soft robot. Using the PWM method, we generated temporal patterns coding that negative pressure was sequentially applied from the posterior to anterior chambers (Figure 7). As an initial case, we set the vacuum pressure as –10 kPa, segmental contraction duration as one second, and overlap of the contraction time between neighbouring segments as zero (Figure 7A). Under this condition, the soft robot exhibited a crawling motion and could move forward (Figure 7B). Accordingly, this result suggests that our soft robot could provide a physical model to analyze the crawling behaviour. In the following sections, we attempted to reproduce two previous experimental results and make one new prediction on fly larval locomotion with the soft robot.

**Figure 7.**
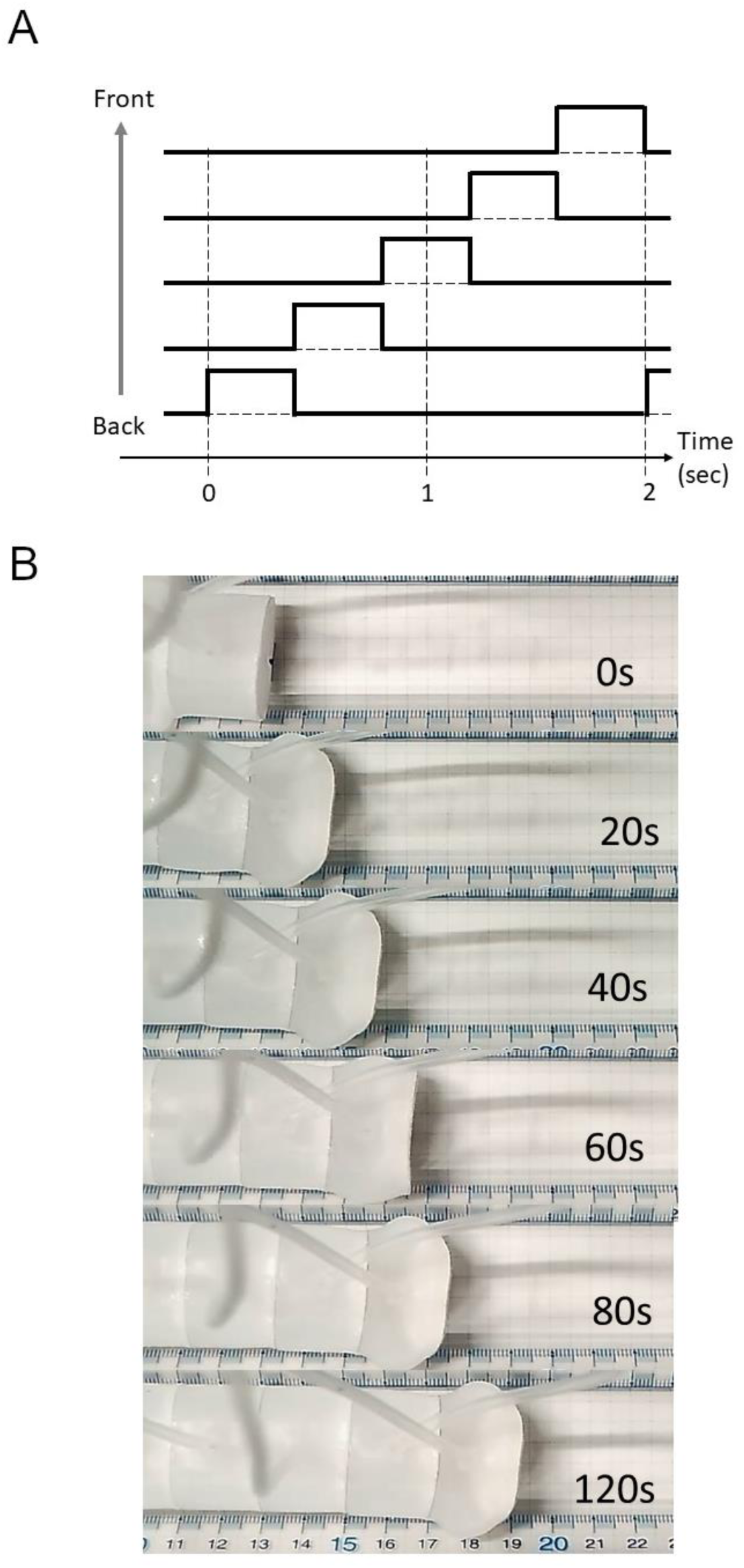
Performance of the soft robots.

### 4.3 Asymmetric speed between forward and backward crawling

As the first experimental observation in fly larvae, we focused on the difference between forward and backward crawling speed. A previous study showed that fly larvae move faster by forward crawling than backward [24]. However, the possible contribution of the difference in the body-ground friction during forward and backward motion to the different speeds between forward and backward crawling has not been tested. By using our soft robot, we tested this possibility. The displacements of the soft robot were calculated based on the position of its front end (the head marker in Figure 8). The speed of robot crawling was measured using five successive strides. To generate backward crawling, we controlled the spatiotemporal pattern of the chamber pressures in a similar way to that of forward crawling but reverse order (Figure 8). We scanned the maximum pressures, stride durations, and intersegmental delays and compared the speed of crawling between backward and forward locomotion. In all the cases, backward crawling was slower than forward crawling (Figure 8C). This observation implies that the asymmetric properties of the interface between the larval body and the ground substrate could be a critical factor in the asymmetric crawling speed between forward and backward movements.

**Figure 8.**
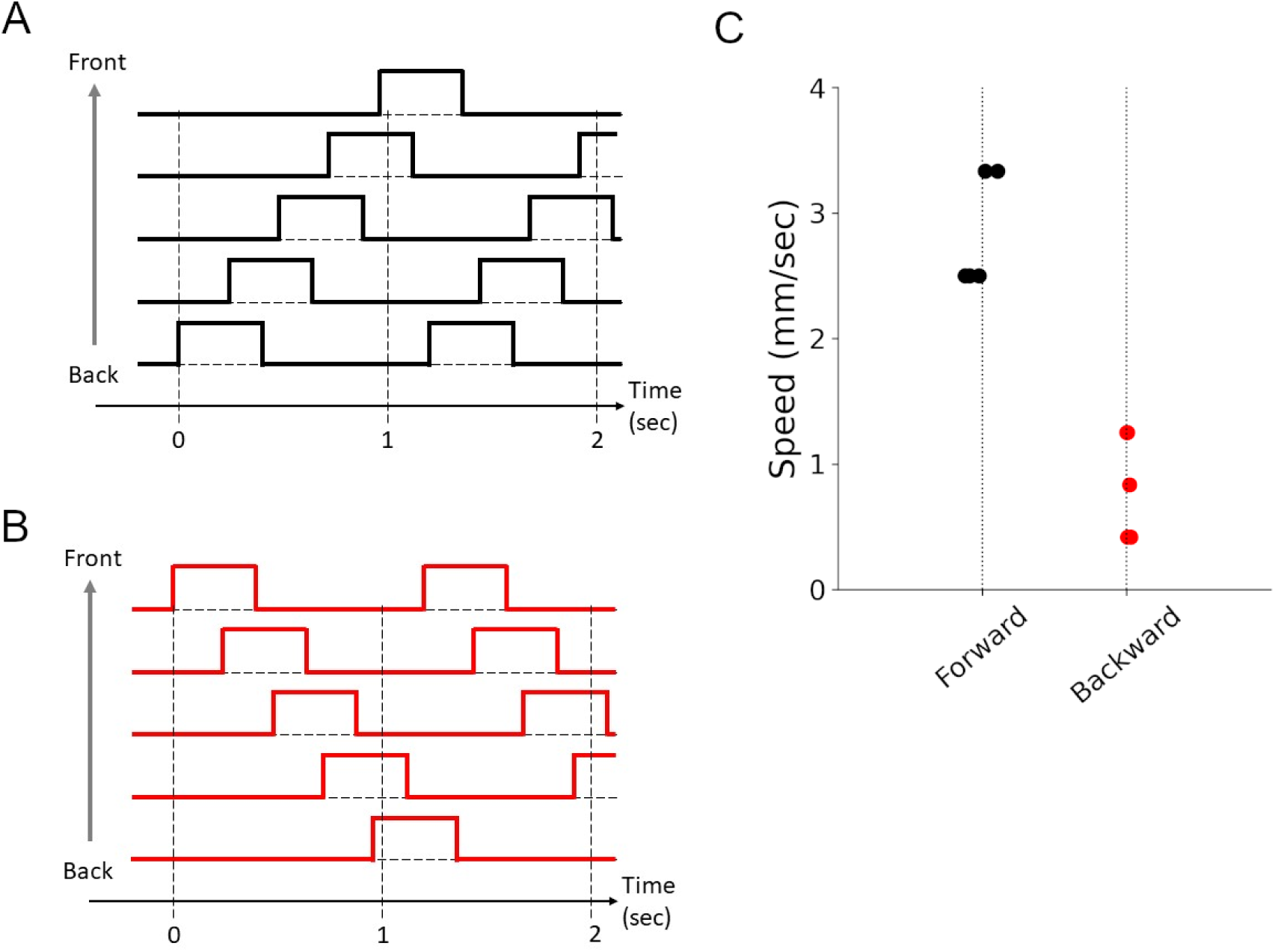
Forward and backward crawling in the soft robot.

### 4.4 Involvement of segmental contraction duration and intersegmental phase delay in locomotion speed

We next analyzed the relationship between the segmental kinematics and crawling speed. A previous study reported that a class of inhibitory interneurons (PMSIs, period-positive median segmental interneurons) was involved in crawling speed in fly larvae [25]. When blocking the activity of PMSIs, the segmental contraction duration and the delay in contraction between neighbouring segments were elongated. Furthermore, the larvae with reduced PMSI activity exhibited slower crawling. These observations suggested that either the segmental contraction duration or the intersegmental phase delay should be involved in crawling speed, but this hypothesis has not been tested. Taking advantage of our soft robot, we investigated this possibility.

First, we changed the segmental contraction duration while keeping the pressure constant and no overlapping between neighbouring segments (Figures 9A and 9B). The result showed that the soft robot with a shorter segmental contraction duration (up to 0.2 seconds) exhibited faster crawling (Figure 9C). This observation was consistent with the observation in the loss of function experiment of PMSIs.

**Figure 9.**
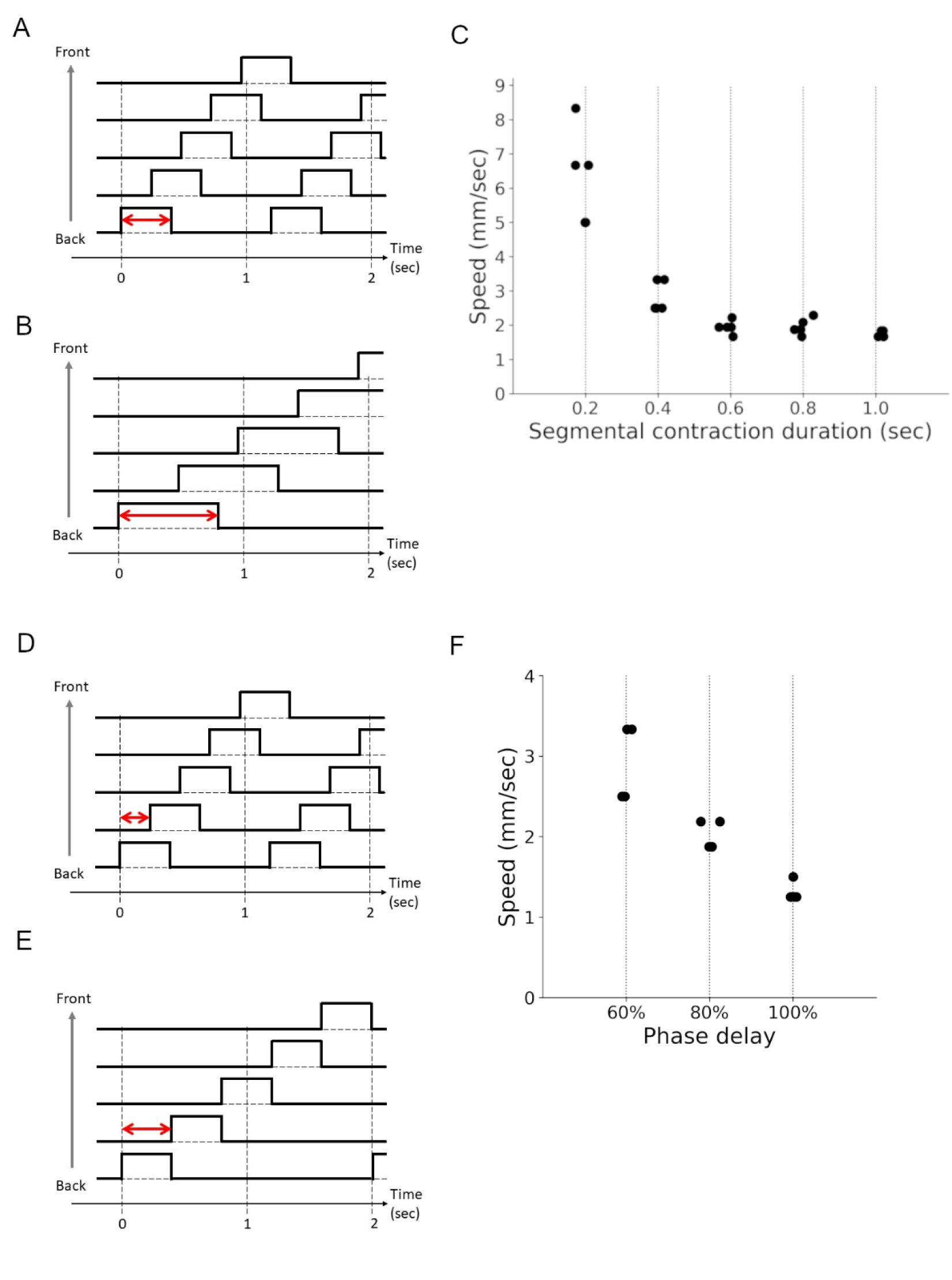
Involvement of intersegmental phase delay and segmental contraction duration in crawling speed.

Next, we perturbed the intersegmental phase delay while keeping other conditions. We defined intersegmental phase delay as the ratio of an intersegmental time delay to segmental contraction duration (Figures 9D and 9E). In fly larvae, the contraction of neighbouring segments overlaps during larval crawling [26]. The stride duration we measured from fly larvae is about one second, and the average segmental contraction duration is around 0.5 seconds. Intersegmental delay is obtained by dividing the stride duration (1 second) by the number of segments (10), giving a 0.1-second intersegmental delay.

Accordingly, the phase delay of neighbouring segments in fly larvae is 20% (= 0.1 sec / 0.5 sec). We investigated the effects of segmental phase delay on crawling speed, which has not been examined before.

We measured robot locomotion with various segmental phase delays. For our five-segment robot, 20% and 40% phase delays were too short to generate stable locomotion because all the segments were shrunk under these conditions. To make time for segments relaxed, we tested segmental phase delays of 60%, 80%, and 100% while keeping segmental contraction time constant (from 0.2 to 1 second) (Figure 9F). We found that as the segmental phase delay increased, the crawling speed became slower (Figure 9F). A similar observation was obtained in operation with different segmental contraction durations (Supplementary Figure 6). These phenomena might be because the smaller segmental phase delay promoted cooperative contraction of neighbouring segments, leading to larger contraction of segments and faster crawling speed (Figure 9). To sum, the perturbation experiments with our soft robot showed that both the segmental contraction duration and the intersegmental phase delay could be involved in crawling speed, consistent with the previous experimental observation.

### 4.5 Prediction of the relationship between the maximum contraction force and locomotion speed

Finally, we analyzed the relationship between segmental contraction force and the crawling speed, which has not been examined in fly larvae yet. We systematically changed the vacuum pressure from –10 kPa to –70 kPa while keeping the segmental contraction duration constant (one second corresponding to the segmental contraction time of 0.2 seconds for a five-segment soft robot) and no overlapping between neighbouring segmental contractions. As a result, the soft robot operated with larger absolute values of the negative pressure and exhibited a faster crawling speed (Figure 10). This tendency could be observed in different segmental contraction durations. This observation could provide a prediction that a larger muscular force could generate faster crawling in fly larvae.

**Figure 10.**
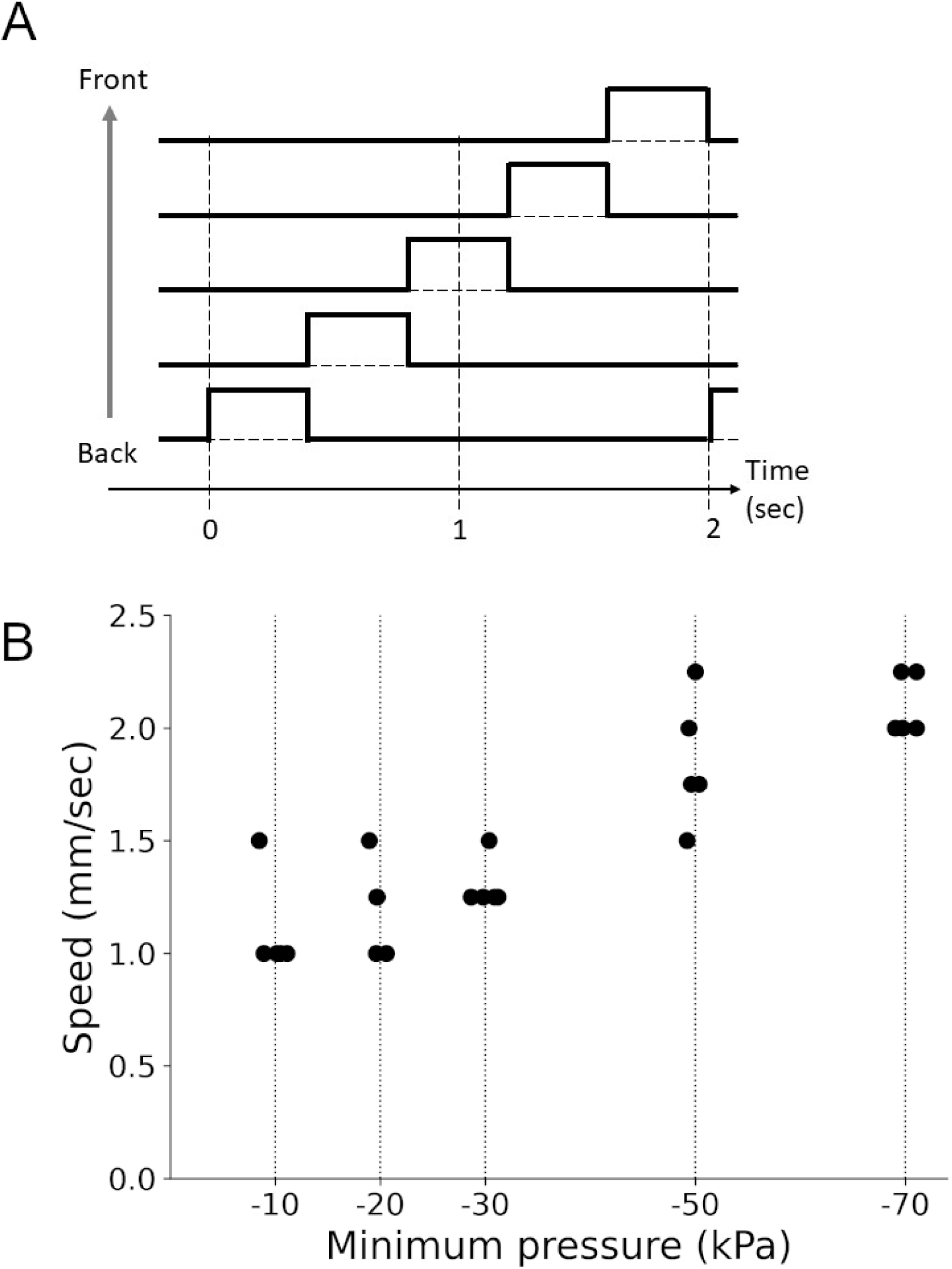
Relationship between the contraction force and crawling speed.

## 5. Discussion

In this work, we developed a new vacuum-actuated soft robot designed to mimic fly larval locomotion. Our soft robot can show peristaltic locomotion patterns produced by the propagation of segmental contraction waves. Two points are crucial to realizing effective peristaltic locomotion in our soft robots: First, pieces of paper inserted in the cross-section help generate asymmetric friction between the soft robot and ground substrate, resulting in different locomotion speeds in different directions. Currently, designs for asymmetric friction in soft robots are still paid little attention. The simple but effective implementation using paper in our robot implies the importance of friction asymmetricity. Second, to determine the control signals for the soft robot, we used the properties in fly larval crawling and configured various parameters, including pressures, stride durations, and intersegmental phase delays via PWM control. Based on the robotic locomotion, we found that the contraction force, intersegmental phase delay, and stride duration all contribute to the regulation of crawling speed.

Analyses using our soft robots are consistent and give novel interpretations to previous works. The mechanism of PMSI neurons indicates that shorter intersegmental phase delays promote faster speed. Assays using soft robots provided evidence (Figure 10D-G) consistent with this. Furthermore, the observation that our soft robot crawls faster in the forward than backward direction is consistent with the previous kinematic study [24]. According to the present study, the difference in speed between forward and backward crawling is partially attributed to the asymmetric friction property between forward and backward directions. Since two separate neural circuits are involved in forward and backward crawling in the central nervous system [18], the different forward and backward crawling speeds might be realized by two parallel mechanisms: friction asymmetricity and circuit specialization.

There are several ways to improve the performance of robotic crawling in future studies. First, the physical properties of soft materials are critical for the dynamics of soft robots. The crawling speed should be improved by utilizing soft materials with larger frictional coefficients for the soft robot. Second, a better actuation source, which can make the soft robot untethered, would broaden its applications in practical scenarios. Third, implementing soft sensors on the robotic body can make it more bionic. It has been reported that the proprioception of the body wall is key to generating an innate crawling speed in *Drosophila* larvae [27]. By monitoring the deformation on the surface of the soft robot and using this information to control the crawling behaviour, the soft robot could exhibit flexible and adaptive locomotion in various complicated environments. Last but not least, although we tested three different sizes of robots, the scale of the robot can still be modified. By changing the number and size of the segments, locomotion ability could be changed. By optimizing these conditions, the application range of our soft larval robots would be broadened.

The kinematic results in our soft robot system have the potential to inspire further study of larval motor outputs, including the mechanisms and kinematic effects of segmental contraction force and phase delay. And our attempts to better understand the mechanisms in soft-bodied animals will contribute to designing adaptive and robust soft robots.

## Acknowledgements

This work was supported by MEXT/JSPS KAKENHI grants (17K19439, 19H04742, and 20H05048 to A.N. and 17K07042 and 20K06908 to H.K.). We thank Dr Jane Loveless and Dr Dai Owaki for their critical comments on our manuscript.

## Conflict of interest

We have no conflict of interest with respect to the work.

## Notes

### Competing Interest Statement

The authors have declared no competing interest.

